# Learning lexical-syntactic biases: An fMRI study on how we connect words and syntactic information

**DOI:** 10.1101/653147

**Authors:** K. Weber, A. Meyer, P. Hagoort

## Abstract

Language processing often involves learning new words and how they relate to each other. These relations are realized through syntactic information connected to a word, e.g. a word can be verb or a noun, or both, like the word ‘run’. In a behavioral and an fMRI task we showed that words and their syntactic properties, i.e. lexical items which were either syntactically ambiguous or unambiguous, can be learned through the probabilities of co-occurrence in an exposure session and subsequently used in a production task. Novel words were processed within regions of the language network (left inferior frontal and posterior middle temporal gyrus) and more syntactic options led to higher activations herein, even when the words were shown in isolation, suggesting combined lexical-syntactic representation. When words were shown in untrained grammatical contexts, activation in left inferior frontal cortex increased. This might reflect competition between the newly learned representation and the presented information. The results elucidate the lexical nature of the neural representations of lexical-syntactic information within the language network and the specific role of the left inferior frontal cortex in unification of the novel words with the surrounding context.

---

In order to produce and understand sentences, we must know the meanings and the grammatical rules of the language. Part of our grammatical knowledge is tightly linked to knowledge about individual words. We know, for instance, that the word ‘cook’ can be either a noun or a verb, but that the semantically similar word ‘chef’ can only be a noun. Similarly, ‘to buy’ and ‘to acquire’ have closely related meanings but only ‘to buy’ can be used both in a double object construction, as in ‘The girl bought the boy a flower.” and with a prepositional object, as in “The girl bought a flower for the boy.”. Thus, we must have learned, through exposure to the languages, which types of syntactic options go with which words. This holds for all kinds of lexical-syntactic information, from word category to verb argument structure information. In this study we investigated how this type of lexically-bound syntactic information is learned, stored, and processed in the brain. Specifically, we looked at the differential contributions of core regions of the language network, left inferior frontal cortex and left middle temporal gyrus, to these processes.

When processing words in context we activate a large left-dominant fronto-temporal network (Friederici and Gierhan 2013; Hagoort 2014; Hagoort and Indefrey 2014). Some models of language processing, such as the Memory, Unification and Control model (Hagoort 2005, 2013), make specific predictions about how lexical-syntactic information might be represented and processed within this language network. The assumption is that syntactic information, especially lexically-bound syntactic information, is stored with the lexical items in the ‘mental lexicon’ (Levelt 1992), located in posterior temporal regions of the brain. When words are processed in context, the left inferior frontal cortex works together with the temporal brain regions to unify the information, selecting the relevant information, and integrating it into context (Hagoort 2005, 2013). These models are in line with lexicalist accounts of language processing, where syntactic information is stored with the lexical items in long-term memory and retrieved in order to be unified into larger linguistic representations (MacDonald et al. 1994; Jackendoff 2002; Culicover and Jackendoff 2005). These models can be investigated by studying lexical-syntactic information. For example, if a word has several, alternating syntactic options this means that it is ambiguous with regards to its syntactic information (Shapiro et al. 1987). In context the ambiguity is usually temporal and the surrounding lexical, syntactic, and semantic information disambiguate between the options. This type of ambiguous lexical-syntactic information, in comparison with unambiguous information, can be used to study the nature of lexical-syntactic representations and processing in the brain.

Several fMRI studies have compared the processing of ambiguous versus unambiguous words, either presented in context or in isolation. Generally, regions of the fronto-temporal language network were more active for syntactically or semantically ambiguous sentences compared to those with low ambiguity (Rodd et al. 2005, 2010). This fronto-temporal language network also showed an increase in activation with the number of syntactic options of a verb, indicating that this syntactic information is stored with the verbs (Shetreet et al. 2006). When presented in isolation, without a sentential context, verbs with multiple compared to single syntactic options showed more activation in angular and supramarginal gyrus as well as superior and middle temporal gyrus (Meltzer-Asscher et al. 2013) supporting the idea of access to lexical-syntactic representations in the posterior temporal and parietal regions of the brain. Also middle and superior frontal gyri showed an ambiguity effect, which the authors related to the processing of the ambiguities themselves.

One study set out to differentiate between the roles of the left inferior frontal and the left posterior middle temporal gyrus directly. The study manipulated both retrieval from the mental lexicon and unification load using a verb-noun category ambiguity paradigm (Snijders et al. 2009). As predicted by the authors and in line with Hagoort (2005, 2013), the left inferior frontal cortex was sensitive to manipulations of unification load, when ambiguous words had to be integrated into a sentence context, but not when the words appeared within a word list. In contrast, the left posterior middle temporal gyrus showed an ambiguity effect both in sentence and word list contexts. This is effect was larger in a sentence context. Retrieving words with two syntactic options compared to one led to more activation independent of context, linked to the retrieval of the lexical-syntactic information from the mental lexicon. Thus, there is evidence for some regional specificity in the brain with regards to storing and processing lexical-syntactic information. This gives some initial insights into the roles and functions of the key regions within the language network that we want to further investigate here.

## Hypotheses

In this study we investigated how novel lexical items and the attached syntactic information are represented and processed within the core language regions of the brain, the left inferior frontal cortex and left posterior middle temporal gyrus. We are interested in investigating both memory and processing aspects related to this type of information. We used two types of lexical-syntactic information, verb-argument structure and verb-noun category ambiguity, to be able to draw conclusions that are not specific to one type of lexical-syntactic information..

We had the following specific hypotheses:

1. **Behaviour:** Participants would learn the lexical-syntactic biases from mere exposure. Behaviourally, their sentence production choices would follow the input. Thus, we predicted that a biased lexical item, which was presented with only one syntactic option, would be used in the biased sentence context. An unbiased lexical item, which was presented with two syntactic options, would be used with both sentence contexts.
2. **Brain activation:** After the initial exposure, the lexical-syntactic representations of the novel words would differ between the biased and unbiased conditions. Unbiased words would have two syntactic options represented with the lexical entry, while biased words would have only one. This would be reflected by increased activation for retrieved words from the unbiased condition within the language network of the brain. This increase in activation might be focused to left posterior middle temporal gyrus as this region is particularly linked to a mental lexicon function for syntactic information (Hagoort 2013). When biased words were exposed with the other sentence context, this unexpected combination should lead to a stronger activation in the language network, more specifically left inferior frontal cortex. This would reflect the competition between the expected and the presented frames or the error signal arising from the unexpected combination. Thus, our main aim was to differentiate the functional roles, i.e. unification and memory, of left inferior frontal cortex and left posterior middle temporal gyrus in lexical-syntactic language processing.

Language is inherently complex and multi-dimensional. It is challenging to investigate memory and unification processes related to the structure of the language independently from semantic aspects. For example, imaginability will covary with number of syntactic frames and the frequencies of occurrence of different verbs. Hence, different syntactic options associated with the verbs make it hard to control for other influencing factors. Therefore, we decided to use a language context with only limited semantic content and to use novel lexical items to have control over input frequencies and the biases people are exposed to. We know from previous research that lexical-syntactic information, such as verb biases, can be quickly learned through exposure via statistical learning (Wonnacott et al. 2008) and this newly learned information is used in subsequent language production (Thothathiri and Rattinger 2015; Thothathiri et al. 2017). What the current study adds is that we used this ability to quickly learn novel lexical-syntactic information to study memory and unification operations during language processing in the brain.

## Materials and Methods

The study included a behavioural experiment (Experiment 1) aiming to establish the participants’ sensitivity to the syntactic biases present in the materials and the main experiment (Experiment 2), aiming to investigate memory and unification processes on novel lexical-syntactic information. Each experiment involved multiple sessions, spread out over two days (see Figure 1).

**Figure 1.**
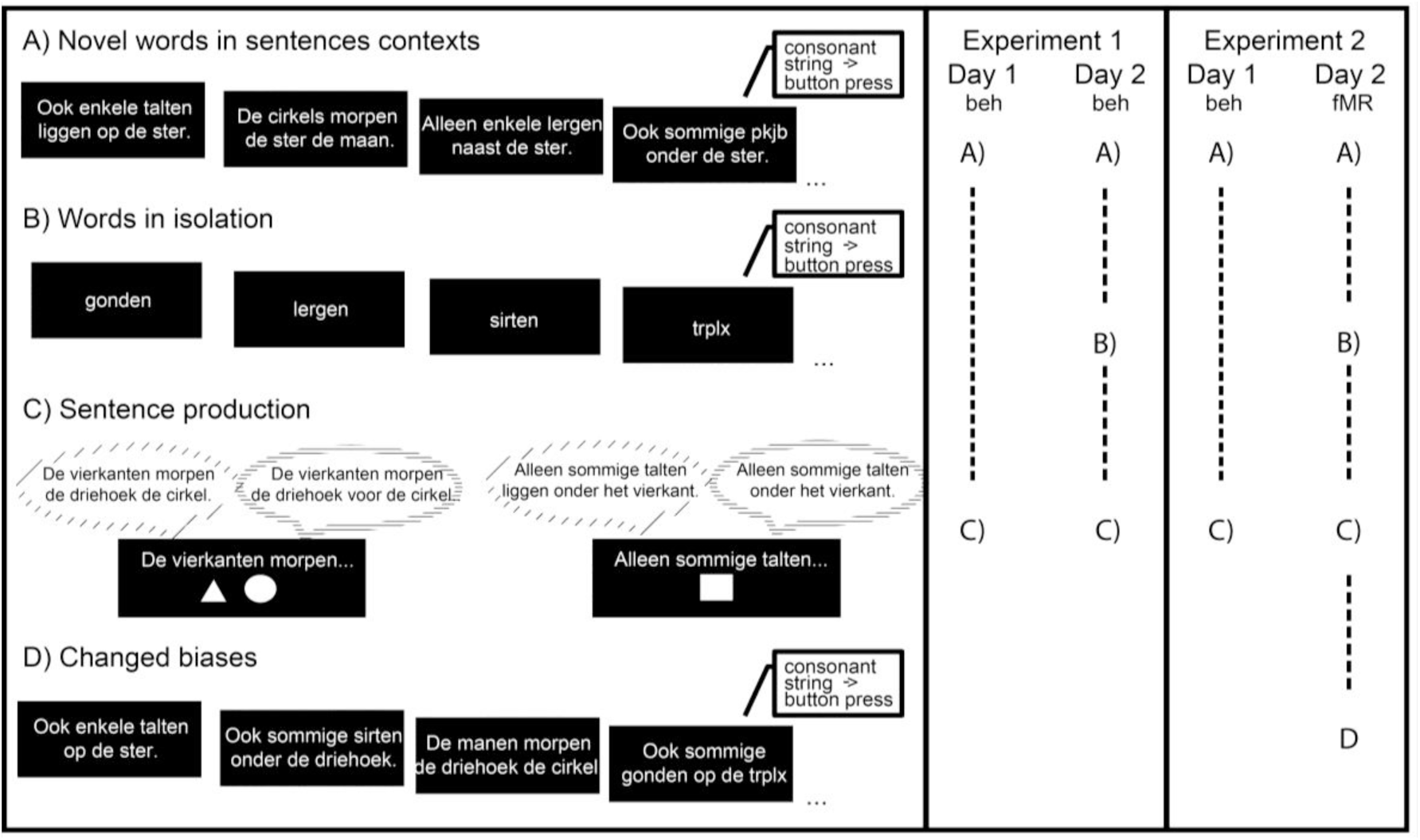
Experimental Design of the four different sessions: A) Participants read the novel words in sentence contexts. B) Participants read the novel words in isolation. C) Participants were presented with the beginning of a sentence and objects and asked to produce a sentence. D) In the last session participants read the novel words in sentence contexts but for half of the biased verbs the biases were changed to the other sentence context. The columns on the right indicate which sessions were run for which experiment and on which day.

### Experiment 1 (behavioural-only)

#### Participants

In the behavioural-only study we tested 32 Dutch native speakers (21 female, 11 male). One additional participant was excluded from the analysis as they frequently used non-learned constructions in the production output. All participants had normal or corrected to normal vision. They were paid for their participation. The participants gave written informed consent prior to participating. The study was conducted according to the institutional guidelines of the local ethics committees (protocol ECG2013-1308-120 for the Max Planck Institute).

#### Materials

Materials were in Dutch, the native language of our participants. The nouns in the sentences were inanimate objects (circle, triangle, moon, star, square) to reduce any potential semantic biases created by the words. The novel words were 12 Dutch pseudowords. They followed the standard phonotactic patterns of Dutch while not having any known semantic meaning, as verified by several native speakers. Sentence contexts were two types of sentences per type of lexical bias category. (1) For the verb-argument structure ambiguity sentence contexts were either prepositional object (Dutch example: “De vierkanten gonden de maan voor de cirkel.”, English literal translation: “The squares gond the moon for the circle.”) or double-object ditransitive sentences (Dutch example: “De sterren welmen de driehoek de cirkel.”, English literal translation: “The stars welm the triangle the circle”). (2) For the sentence contexts with a verb-noun category ambiguity, one context ensured a noun reading of the novel word (Dutch example: “Alleen sommige talten zitten onder het vierkant” English literal translation: “Only some talts sit under the square”) while the other type of context ensured a verb reading (Dutch example:”Alleen sommige talten onder het vierkant”, English literal translation: “Only some talt under the square”). A full list of the novel words and types of sentence contexts can be found in Appendix A. There were sets of two verbs per condition (randomly assigned to the conditions; 4 different lists across participants) combined with random assignments of the 5 nouns (no noun was repeated within a sentence context).

#### Design and Procedure

The experiment was run over two days with different experimental sessions:

A. *Novel words in sentence context:* In these sessions participants read the novel words embedded in sentence contexts (Figure 1, A). The factors were ‘type of bias’ (verb-argument structure or verb-noun category ambiguity) and ‘bias’ (100% bias 1; 50% bias 1/50% bias 2; 100% bias 2). To ensure attention some sentences included a word made up of consonants to which participants were asked to press a button. Sentences were displayed on the screen for 3.5 seconds with a variable inter-trial interval between 1 and 3 seconds (to mirror the jitter needed in the MRI experiment).
B. *Words in isolation:* In this session participants read the novel words in isolation (see Figure 1, B). The factors were ‘type of bias’ (verb-argument structure or verb-noun category ambiguity) and ‘bias’ (100% bias 1; 50% bias 1/50% bias 2; 100% bias 2. To ensure attention some items were made up of consonants to which participants were asked to press a button. Words were displayed on the screen for 1.8 seconds with a variable inter-trial interval between 1 and 3 seconds (to mirror the jitter needed in the MRI experiment).
C. *Sentence production:* In the production sessions participants received the initial part of the sentence context until the novel word and were asked to make a full sentence including the one or two objects shown on the screen (see Figure 1, C). The sentence fragment and the objects were displayed on the screen for 4.4 seconds with a variable inter-trial interval between 1 and 3 seconds (to mirror the jitter needed in the MRI experiment).

In all sessions conditions were displayed in random order.

On the first day participants participated in 2 sessions, an ‘novel words in sentence contexts’ (A) session, and a short ‘sentence production’ (C) session. On the second day there was another ‘novel words in sentence contexts’ (A) session, a ‘words in isolation’ (B) session (to mirror the fMRI experiment) followed by a longer ‘sentence production’ (C) session. Session (D) was only run for Experiment 2.

The ‘novel words in sentence contexts’ session included 30 trials per condition on day 1 and 20 trials on day 2. The ‘sentence production’ session consisted of 4 trials on day 1 (to familiarize participants with the task) and 20 trials on day 2. Reading the ‘words in isolation’ was done for 20 trials per condition. Participants sat comfortably in front of a computer monitor. Sentences were displayed in white Calibri font, font size 16 on a black background. In the production session the objects to be included in the sentences were displayed below the sentence fragments.

Participants’ verbal responses during the production session on Day 2 were recorded via a microphone. Trials with incomplete utterances, structures that did not match any of the possible input structures, and those with reaction times 3 standard deviations above or below the mean of the overall reaction times per participant were removed from the analyses.

#### Behavioural Analysis

We analyzed the production output of Day 2 in mixed-effects logit models (Pinheiro and Bates 2000; Jaeger 2008; Barr et al. 2013) using a mixed effects model with random effects for subjects and items in R (*RStudio: Integrated Development for R* 2013) and the lm4 toolbox (Bates et al. 2015). Following (Barr et al. 2013), we chose a model with the maximal effect structure that still converged. When a model did not converge, we removed random slopes for the factors with the lowest variance first. We used a model per ‘type of bias’ (verb-argument structure or verb-noun category ambiguity) with ‘bias’ (100% bias 1; 50% bias 1/50% bias 2; 100% bias 2) as a factor. For contrast specifications, treatment coding was used, where we compared the reference level (the 50% bias 1/50% bias 2 condition) to the other two conditions.

### Experiment 2 (fMRI and behavioural)

#### Participants

In a second experiment we added fMRI scanning as an additional measure. In this experiment we tested 31 Dutch native speakers (22 female, 9 male). 7 additional participants were tested but excluded from the analysis as 2 participants did not complete Day 2, 1 showed excessive movement (up to 15mm), for 1 participant we had technical problems, 1 participant frequently used non-learned constructions in their production output and 2 participants did not perform the behavioural task during the analysed experimental sessions. The participants were all right-handed and had passed screening for MRI compatibility. All participants had normal or corrected to normal vision. The participants received money for their participation. All participants gave written informed consent prior to participating. The study was conducted according to the institutional guidelines of the local ethics committees (CMO2014/288 for the Donders Institute).

#### Materials

The stimulus material was the same as for Experiment 1.

#### Design and Procedure

The experiment was run over two days with similar sessions as the behavioural-only experiment (Experiment 1):

A. Novel words in sentence contexts: see Experiment 1 above
B. Words in isolation: see Experiment 1 above
C. Sentence production: see Experiment 1 above
D. Changed biases: In this session participant read the novel words embedded in sentence contexts but for half of the words that had been displayed with a 100% bias to one sentence context, the context was changed. The factors were ‘type of bias’ (verb-argument structure or verb-noun category ambiguity) and ‘change’ (changed bias; unchanged bias). To ensure attention some sentences included a word made up of consonants to which participants were asked to press a button. Sentences were displayed on the screen for 3.5 seconds with a variable inter-trial interval between 1 and 3 seconds.

In all sessions conditions were displayed in random order. On the first day participants participated in two sessions, that were behavioural-only, a ‘novel words in sentence contexts’ session, and a short ‘sentence production’ session. On the second day all experimental sessions were conducted in the MR scanner. It started with a ‘novel words in sentence contexts’ session, followed by the session in which the novel words were read in isolation, and another production session. This was followed by a ‘changed biases’ session. The ‘novel words in sentence contexts’ session included 32 trials per condition on day 1 and 18 trials on day 2 (with an additional 12 consonant string trials for each day, trial numbers were slightly adapted from the behavioural study as we had to shorten the fMRI scanning session on day 2). The ‘words in isolation session’ consisted of 32 trials per condition (plus 12 task trials). The production session consisted of 2 trials per condition on day 1 (to practice the task) and 32 trials on day 2. The ‘changed biases’ session consisted of 28 trials per condition (plus 10 task trials). Participants sat comfortably in front of a computer monitor. Sentences were displayed in white Calibri font, font size 20 on a black background. In the production session the objects to be included in the sentences were displayed below the sentence fragment. Participants’ verbal responses during the production session on Day 2 were recorded via an MR compatible microphone.

Trials with incomplete utterances, structures that did not match any of the possible input structures and those with reaction times 3 standard deviations above or below the mean of the overall reaction times per subject were removed from the analyses. Preprocessing and analysis of the fMR images was done using SPM 12 (*Statistical Parametric Mapping* 2014).

#### Behavioural Analysis

See behavioural analysis of Experiment 1 above.

#### FMRI data acquisition

MRI data were recorded in a 3 T MR scanner (PrismaFit, Siemens Healthcare, Erlangen, Germany) using a 32-channel head coil. Whole-brain functional images were collected using a multi-band (accelerator factor of 8) T2*-weighted sequence: repetition time (TR): 735 ms; echo time (TE) 39 ms; field of view 210 × 210 mm; 64 slices; voxel size 2.4 × 2.4 × 2.4 mm. To correct for distortions, fieldmap images were also recorded. Additionally, T1-weighted anatomical scans at 1 mm isotropic resolution were acquired with TR 2300 ms, TE 3.03 ms, flip angle 8°, and FOV 256 × 256 × 192 mm.

#### FMRI data preprocessing

First, DICOM images converted to nifti images. Then functional volumes were realigned and unwarped using the acquired fieldmaps, and coregistered to the individual structural image and further normalized to a standard MNI space (resampled at voxel size 2 × 2 × 2 mm). Lastly, the images were spatially smoothed with a kernel of 5 mm full width at half maximum.

#### FMRI analysis

##### First Level – Words in Isolation (B)

For the design matrix for the ‘words in isolation’ part of the study we modelled event-related regressor for each of the conditions of the factor ‘Type of Bias’ and ‘Bias’. The onset of the word was taken as the time of onset and the actual duration of word presentation was modelled. In addition we added a regressor for those trials on which participants had to perform the behavioural task and we also added 6 movement regressors. Per subject we identified contrast images that were then taken to the second level for a random effects group analysis.

##### First Level – Changed biases (D)

For the design matrix for the ‘changed biases’ part of the study we modelled event-related regressor for each of the conditions of the factor ‘Type of Bias’ and ‘Novelty’. The onset of the sentence was taken as the time of onset and the actual duration of sentence presentation was modelled. In addition we added a regressor for those trials on which participants had to perform the behavioural task and we also added 6 movement regressors. Per subject we identified contrast images that were then taken to the second level for a random effects group analysis.

##### Second Level Analysis

As we were interested in two specific regions of interest (the left inferior frontal cortex and the left posterior middle temporal gyrus) we focused our group analysis on these. The regions were defined based on the left inferior frontal and posterior middle temporal activations for the term ‘syntactic’ with the meta-analysis toolbox neurosynth (Yarkoni et al. 2011) at a threshold of Z>6. Mean contrast values per region, condition and subject were extracted using MarsBar (Brett et al. 2002) and analysed using an ANOVA in SPSS (*IBM SPSS Statistics for Windows* 2011). Additional whole-brain analyses are described in the Supplementary Materials (Appendix B).

## Results

### Experiment 1 – behavioural results

The model of the production output included fixed effects for ‘Bias’ (100% bias 1; 50% bias 1/50% bias 2; 100% bias 2). The random effects structure included a random intercept for subjects and items and random slopes for ‘Bias’ for subjects and items.

The participants showed the expected staircase pattern (see Figure 2 for a figure of the group averages and Supplementary Figure 2 for individual subject data) with the 50% bias 1/50% bias 2 conditions patterning in the middle. In the verb-argument set the PO structure was used more often than the DO structure. In the verb-noun category ambiguity set the word was most often used as a verb. For the model investigating the verb argument structure manipulation, the contrast reflecting differences between DO bias condition and the 50% bias 1/50% bias 2 condition was significant (β=-1.26, SE=.43, Z=-3.0, p=.003) as was the other contrast between the PO condition and the 50% bias 1/50% bias 2 condition (β=.86, SE=.37, Z=2.3, p=.022). For the verb-noun category ambiguity manipulation, the contrast between the noun condition and the 50% bias 1/50% bias 2 condition was not significant (Z=1.1), while the contrast between the verb and the 50% bias 1/50% bias 2 condition was (β=-1.3, SE=.49, Z=-2.65, p=.008).

**Figure 2.**
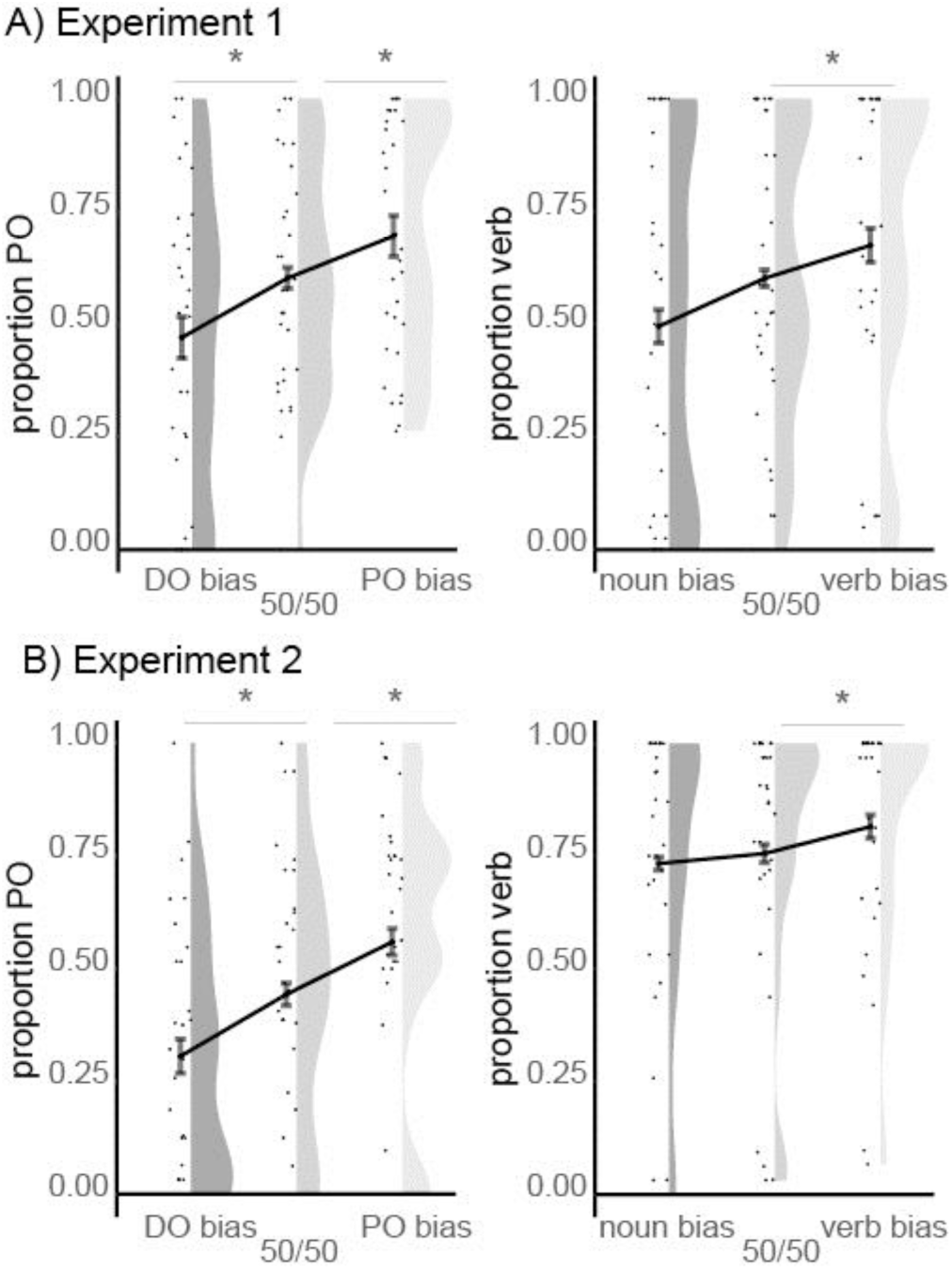
Behavioural results – Production choices in the behavioural-only experiment 1 (top) and fMRI experiment 2 (bottom). Left: Proportion of PO structure sentences used in production output per condition for the verb-argument structure manipulation. Right: Proportion of verb usage in production output per condition for the verb-noun category manipulation. The graphs plot individual data point, the mean and standard error of the mean on the left and density on the right.

### Experiment 2 – behavioural results

The model of the production output included fixed effects for ‘Bias’ (100% bias 1; 50% bias 1/50% bias 2; 100% bias 2). The random effects structure included a random intercept for subjects and items and random slopes for ‘Bias’ for subjects and items.

The participants showed the expected staircase pattern (see Figure 2 for a figure of the group averages and Supplementary Figure 2 for individual subject data) with the 50% bias 1/50% bias 2 conditions patterning in the middle.

For the verb argument structure manipulation the DO structure was used more often overall and for the verb-noun category ambiguity manipulation the word was most often used as a verb. For the model investigating the verb argument structure manipulation, the contrast reflecting differences between DO bias condition and the 50% bias 1/50% bias 2 condition was significant (β=-.84, SE=.34, Z=-2.4, p=.015) as was the other contrast between the PO condition and the 50% bias 1/50% bias 2 condition (β=.81, SE=.21, Z=3.85, p<.001). For the verb-noun category ambiguity manipulation, the contrast between the noun condition and the 50% bias 1/50% bias 2 condition was not significant (Z<1), while the contrast between the verb and the 50% bias 1/50% bias 2 condition was (β=.97, SE=.47, Z=2.07, p=.04).

### Experiment 2 – fMRI results

#### ROI analysis - Words in isolation

Overall there was a main effect of bias, F(1,30)=7.53, p=.01 as the words with two syntactic options (unbiased) showed more activation than the ones with only one option (biased), see Figure 3A for an illustration. None of the other main effects or interactions reached significance. Specifically, there was no interaction between bias and region (F<1) or between bias, type of bias and region showing that the effect is not specific to a particular region in the language network. Follow-up one-tailed t-tests were run to investigate the effect of unbiased>biased for each ‘Type of Bias’ and region of interest. For these 4 tests the alpha level was adjusted to .0125. In LIFC, the bias effect for the verb-argument structure was marginally significant (t(30)=1.69, p=.05) and significant for the verb-noun category ambiguity condition (t(30)=2.83, p=.004). In LMTG, the effect was not significant for the verb-argument structure condition, |t|<1 but significant for the verb-noun category ambiguity condition (t(30)=2.68, p=.006).

**Figure 3.**
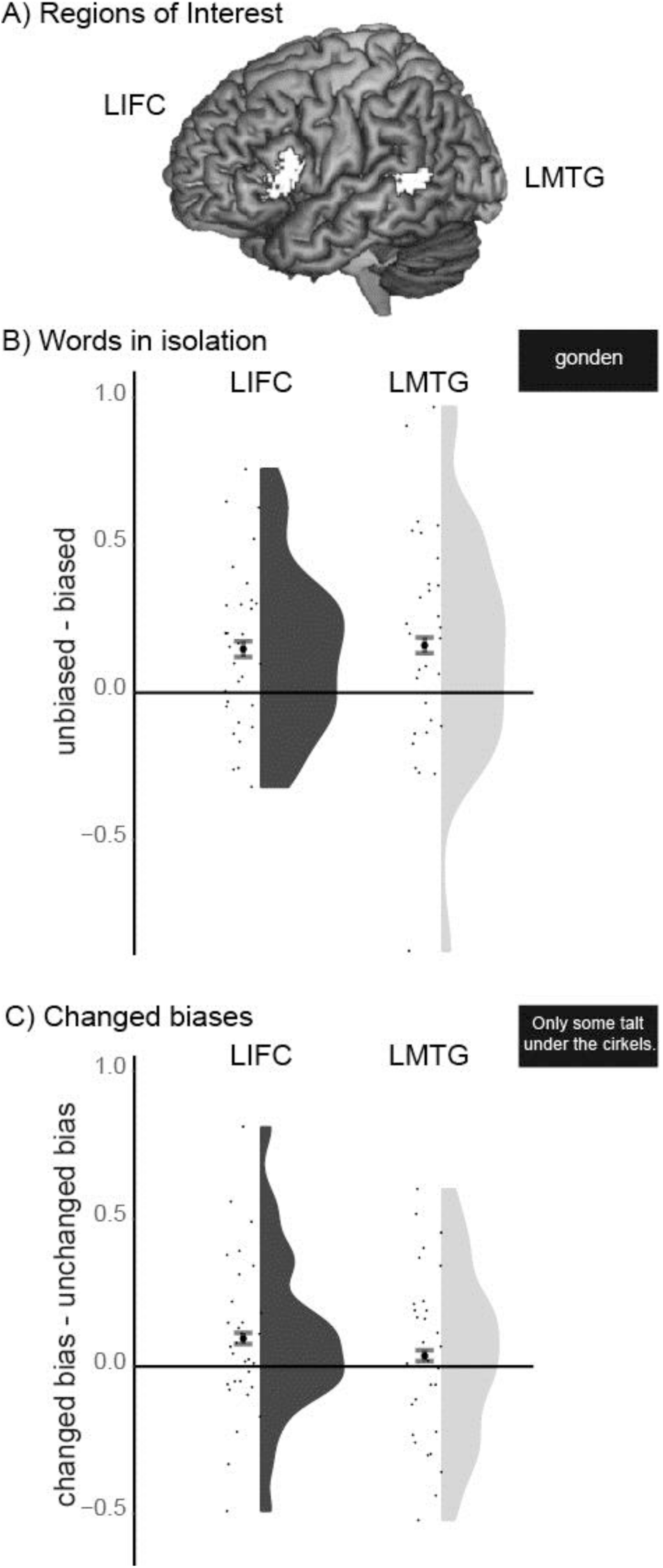
Region of interest results. A) The regions of interest in left inferior frontal cortex (LIFC) and left middle temporal gyrus (LMTG). B) Reading the novel words in isolation led to enhanced activation for novel words for which two syntactic options were given in the sentence exposure session (unbiased) compared to those with only one syntactic option (biased). This effect of ‘bias’ is significant over both regions of interest. C) In LIFC those sentences where the novel words were presented in changed sentence context, with which they were not paired in the initial sentence exposure session, showed higher activation than those with the old sentence contexts. The graphs plot individual data point, the mean and standard error of the mean on the left and density on the right.

A whole-brain analysis (see Supplementary Materials, Appendix B), using a flexible factorial design with the factors ‘type of bias’ and ‘novelty’, showed mainly overlapping results.

#### ROI analysis - Changed biases

In the ‘changed biases’ part of the study, bias change (changed bias; unchanged bias) interacted with region (LIFC; LMTG), F(1,30)=4.8, p=.036. As predicted, in LIFC a changed bias led to higher activation than the unchanged bias (t(30)=2.03, p(1-tailed)=.026), while in LMTG it did not, |t|<1 (see Figure 3C). None of the other main effects or interactions reached significance. A whole-brain analysis (see Supplementary Materials, Appendix B), using a flexible factorial design with the factors ‘type of bias’ and ‘novelty’, did not show any significant effects.

## Discussion

### Behavioural production choices

From the behavioural data in both experiments it is evident that participants learned the lexical-syntactic biases very quickly through mere exposure. As expected, the production choices followed the probabilistic nature of the input. During sentence production participants were more likely to use the novel lexical items with the syntactic contexts they were paired with during the learning phase. This was the case for both types of lexical-syntactic information, verb argument structure as well as category ambiguity biases. This is in line with previous studies showing a similar pattern for verb-argument structures only (Wonnacott et al. 2008; Thothathiri et al. 2017). This was mostly likely the case because the syntactic options for each novel lexical item were quickly integrated into the lexical item’s memory representation. This is further supported by the neuroimaging findings.

### Neural effects of the novel lexical-syntactic representations

The first part of the fMRI experiment investigated the nature of the newly learned lexical-syntactic representations and their location in the brain when the words were presented in isolation after the learning phase. The region of interest analysis in LIFG and LMTG revealed that the syntactic options are combined with the lexical item within the language network. Even without a context, lexically-syntactic ambiguous words, those with two syntactic options, showed an increase in the hemodynamic response in comparison to unambiguous words. This pattern indicates that item-specific syntactic options were stored with the item within the language network, leading to higher activation when a lexical item with two syntactic options was accessed in memory. While we had predicted this effect to occur in left posterior temporal gyrus specifically, in line with the memory part of the MUC model (Hagoort 2005, 2013), it was present in the left posterior temporal gyrus as well as the left inferior frontal cortex. This finding could be indicative of a more distributed representation of lexical items and the lexical-syntactic options within the language system, especially during learning. However, it could also be the case that, given the recency of the exposure in context, participants were attempting to build relevant contexts online. This could thus potentially still be in line with a model that posits the location of the lexical-syntactic representation to be in posterior temporal regions with the left inferior frontal regions related to processing. In the whole brain analysis, the left inferior frontal effect extends into the left insular cortex. While this is not one of the regions we had proposed for the contrast of two versus one syntactic options when reading words in isolation, it is a region that has been found in other studies on ambiguity processing in general, such as ambiguous meanings (Mason and Just 2007).

Interestingly, the pattern of activation did not differ between the types of lexical-syntactic information, verb-argument structure and verb-noun category ambiguity. This thus speaks to the general nature of lexical-syntactic representations in the brain, where at least some syntactic information is stored in the lexicon.

### The role of LIFC

Within this experiment we were also able to show a more specialized function of left inferior frontal cortex. In contrast to LMTG, we found a unique contribution of LIFC when biased words were exposed with the other sentence context. Thus, the unexpected lexical-syntax combination, as predicted, led to stronger activation in left inferior frontal cortex specifically than a learned lexical-syntax combination.

This effect might reflect competition between potential frames or category assignments driven by lexical-syntactic information (Vosse and Kempen 2000). This proposal is in line with the MUC model (Hagoort 2005, 2013) that posits that the building blocks from memory are unified through the involvement of the left inferior frontal cortex. During this process appropriate items have to be selected and alternatives inhibited. In the model by Vosse and Kempen (2000) this is done by lateral inhibition between alternative candidates. Our findings go beyond the earlier finding of a larger ambiguity effect in sentence contexts in LIFC reported by Snijders and colleagues (2009), as in their study the effect was not shown to be significantly different from the effect in LMTG. Here, we show a clear specification of LIFC in comparison to LMTG for unification related operations, in line with the MUC model (Hagoort 2005, 2013).

Another explanation of the increased activation to unexpected lexical-syntactic combinations is that the item-specific lexical-syntactic information leads to expectations regarding the structure a word should appear in. This is in line with computational models of language processing (Chang et al. 2006, 2012). If the predictions based on the lexical-syntactic information are not met a prediction error, or surprisal effect, due to the presentation of an unexpected structure leads to increased activation in left inferior frontal cortex, which might subsequently propagate to the lexical representations in LMTG. Surprisal effects related to syntactic information have previously been found to lead to an increase in activation in left inferior frontal cortex, among other language regions (Weber et al. in press; Bonhage et al. 2015; Henderson et al. 2016).

## Conclusion

In our study, participants adapted to the statistics of the linguistic input, even to subtle lexically driven cues to syntactic information, in line with previous findings on the acquisition and use of probabilistic lexical cues (Wonnacott et al. 2008; Snijders et al. 2009; Fine and Jaeger 2011; Ryskin et al. 2017; Thothathiri et al. 2017). The novel words and the related syntactic options were quickly learned, combined, stored and processed in the language network of left inferior frontal and left posterior middle temporal gyrus.

## Acknowledgments

We thank Vera van’t Hoff, Maarten van den Heuvel, Michel-Pierre Jansen and Birgit Knudsen for their help in behavioural data acquisition and experiment and response coding, as well as Paul Gaalman for help with the fMRI scanning.

## Appendix A

### Example stimuli

Nouns used: Triangle (driehoek), square (vierkant), star (ster), moon (maan), circle (cirkel)

Novel lexical items: 1. Sirten; 2. Morpen; 3. Jurfen; 4. Narten; 5. Welmen; 6. Dalsen; 7. Sulpen; 8.

Polken; 9. Talten; 10. Mupsen 11. Gonden; 12. Lergen Verb-argument structure

DO condition: The noun *novel verb* the noun the noun PO condition: The noun *novel verb* the noun for the noun

Verb-noun category

Verb condition: Conjunction quantifier *novel verb* preposition the noun. Noun condition: Conjunction quantifier *novel noun* verb preposition the noun

Known Dutch words were randomly assigned to a sentence while ensuring no word repetition within a sentence.

## Appendix B

### Whole-brain analysis ‘words in isolation’

We built a flexible factorial design with a regressor per experimental condition (Argument Structure – Biased; Argument Structure – Unbiased; Category Ambiguity – Biased; Category Ambiguity – Unbiased) as well as regressors to model the within subject-effect (thus one regressor per subject).

As expected a region in left inferior frontal cortex showed more activation to unbiased words (with 2 syntactic options) than biased words with only one syntactic option (cluster-level pFWE=.001). Also the other expected region, left posterior middle temporal gyrus (x=-64, y=-44, z=6), showed such an effect but it did not survive cluster-level correction (pFWE=.71). Other regions that showed significantly more activation to unbiased than biased words were located in the left and right posterior insula/rolandic operculum and the right anterior superior temporal gyrus.

There no significant activations for the main effect of ‘type of bias’ or the interaction between ‘type of bias’ and ‘bias’.

**Supplementary Figure 1.**
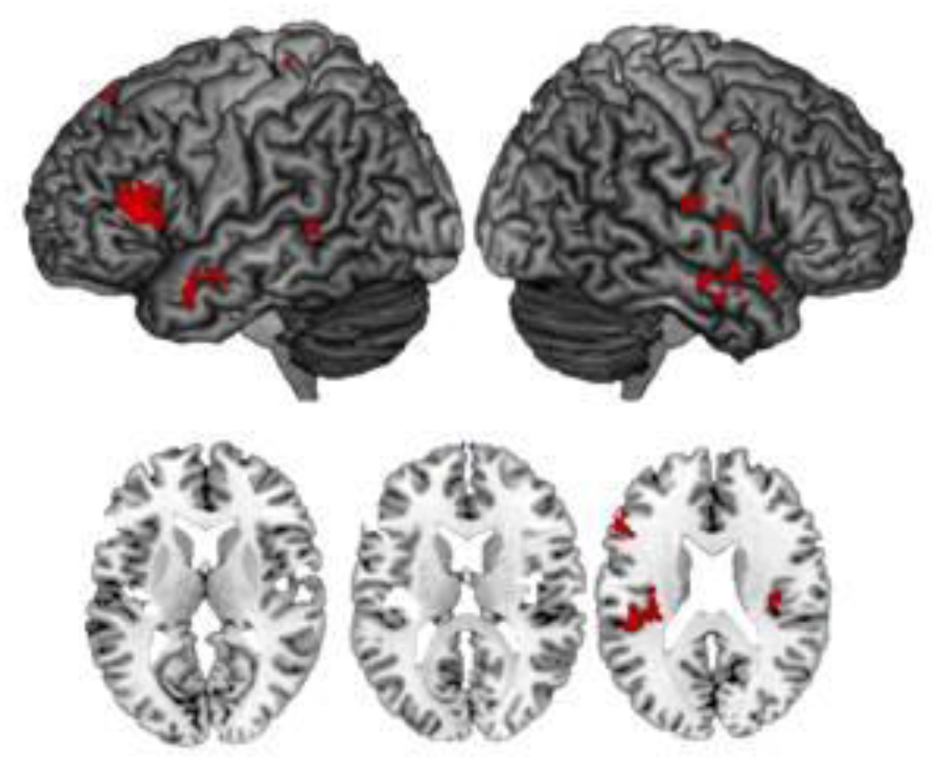
Activations for the main effect of Bias (Unbiased words > Biased words) at a voxel-level threshold of p<.001, k=25. See Table 1 for a list of activations that survive cluster-level family-wise error correction.

**Table 1.** Whole-brain activations for ‘words in isolation’. Listed are local maxima more than 20mm apart. All clusters at a voxel-level threshold of p<.001, and a cluster-level threshold of pFWE<.05 are reported.

| Anatomical label | Global and local maxima |  |  | cluster size k | cluster-level p | Z |
| --- | --- | --- | --- | --- | --- | --- |
|  | x | y | z |  |  |  |
| **Main effect of Bias (Unbiased words>Biased words)** |  |  |  |  |  |  |
| Left insula | -36 | -20 | 14 | 407 | 0.000 | 4.84 |
| Left inferior frontal cortex (triangularis) | -56 | 26 | 12 | 210 | 0.001 | 4.53 |
| Right rolandic operculum | 38 | -18 | 16 | 194 | 0.002 | 4.37 |
| Right superior temporal gyrus | 56 | 0 | -12 | 127 | 0.019 | 4.29 |
| **Main effect of Bias (Biased words> Unbiased words)** |  |  |  |  |  |  |
| - |  |  |  |  |  |  |

### Whole-brain analysis ‘changed biases’

We built a flexible factorial design with a regressor per experimental condition (Argument Structure – Changed Bias; Argument Structure – Unchanged Bias; Category Ambiguity – Changed Bias; Category Ambiguity – Unchanged Bias) as well as regressors to model the within subject-effect (thus one regressor per subject).

There were no significant effects of ‘novelty’ or ‘type of bias’ and no interaction between ‘novelty’ and ‘type of bias’ at a voxel-level threshold of p<.001 and a cluster threshold of pFWE<.05.

## Appendix C

**Supplementary Figure 2.**
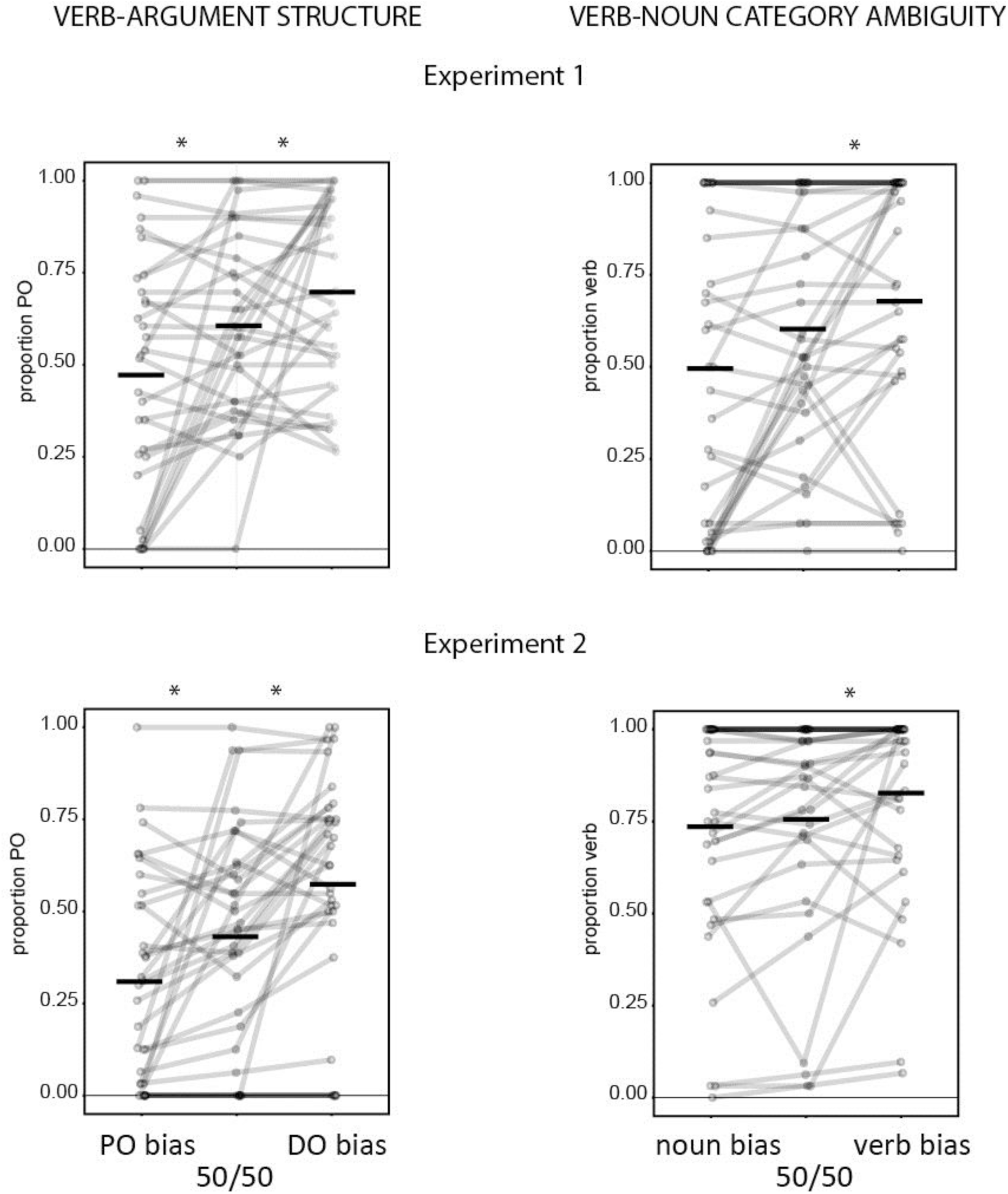
Individual subjects’ production choices in the behavioural-only experiment 1 (top) and the fMRI experiment 2 (bottom). Top Left: Proportion of PO structure sentences used in production output per condition. Top Right: Proportion of verb usage in production output per condition. Bottom Left: Proportion of PO structure used in production output per condition. Bottom Right: Proportion of verb usage in production output per condition.

